# CD47 Blockade Enhances Immunoradiotherapy Response in Head and Neck Squamous Cell Carcinoma

**DOI:** 10.1101/2025.06.29.662015

**Authors:** Abdula Monther, Riyam Al-Msari, Robert Saddawi-Konefka, Santiago Fassardi, Cynthia Tang, Chad Philips, Prakriti Sen, Pardis Mohammadzadeh, Kelsey Decker, Sayuri Miyauchi, Souvick Roy, Riley Jones, Silvio Gutkind, Andrew Sharabi, Joseph Califano

## Abstract

Head and neck squamous cell carcinoma (HNSCC) is a significant cause of morbidity and mortality worldwide, with limited treatment options for patients with locally advanced disease. CD47 immune checkpoint inhibitors have been used to block the CD47/SIRPa interaction that inhibits antigen-presenting cell phagocytosis, thereby enhancing antigen presentation to cytotoxic T-cells, and have shown promise in combination with anti-PD1 immunotherapy in tumors, including recurrent/metastatic HNSCC. We found that CD47 expression is associated with poor prognosis in HNSCC and explored the anti-tumor activity of CD47 blockade in combination with anti-PD1 and lymphatic-sparing radiotherapy in a locally advanced HNSCC model. Using the 4MOSC1 orthotopic, syngeneic murine model of HPV-negative HNSCC, treatment with an engineered CD47-blocking SIRPα fusion protein (ALX301) similarly induced complete tumor regression in combination with anti-PD1, and a partial response as a standalone therapeutic. An anti-PD1 immune checkpoint inhibitor in a CD47-null tumor background led to complete tumor regression confirming a key role for CD47 in tumor immunity. Anti-CD47 treated mice demonstrated increased MHC-II expression on dendritic cells within the tumor and upregulation of CD86 co-stimulatory molecule on dendritic cells within the tumor, sentinel lymph nodes, and contralateral lymph nodes. Combination ALX301 and anti-PD1 treatment in an anti-PD1 resistant 4MOSC2 model demonstrated significant tumor regression, enhanced survivability, improved response with neoadjuvant radiotherapy, and greater retention of CD8+ T-cells within the tumor microenvironment. Notably, T-cell receptor sequencing revealed increased shared clonality between the tumor and sentinel lymph nodes of anti-CD47 treated mice. These data demonstrate that a combination of CD47 blockade and anti-PD1 therapy enhances tumor antigen presentation and immune cell infiltration, while further improving anti-tumor responses in combination with tumor-targeted radiotherapy. This study provides support for the rational design of combinatorial immunoradiotherapy, using anti-CD47 inhibitors and anti-PD1 therapy, in a clinical trial targeting locally advanced HPV-negative HNSCC.

## Introduction

Head and neck squamous cell carcinomas (HNSCC) arise from the upper aerodigestive tract, including the oral cavity, pharynx, larynx, and paranasal sinuses^4,5^. They represent the seventh most common cancer worldwide, accounting for 3% of all cancers, with approximately 890,000 new cases and 450,000 deaths annually^6^. Current therapeutic approaches for HNSCC patients include combinations of surgery, radiation, chemotherapy, immunotherapy, and rarely targeted therapies^7,8^. Despite the advancement in those therapeutic techniques, the overall survival rate has not improved significantly, particularly for patients with locally advanced diseases^7,8^.

Radiotherapy and surgery are primary treatment modalities for patients with locally advanced HNSCC. Therapeutic elective nodal irradiation (ENI) and removal of tumor-draining lymph nodes (neck dissection) are standard of care therapy to reduce regional recurrence in HNSCC^9,10^. Blocking PD-1 interaction with its ligand PD-L1 rescues T cells from exhausted status and revives antitumor immunity^11^. PD-1 inhibitors have shown an overall response rate of 10-20% in recurrent/metastatic HNSCC^12,13^. Recently, PD-1 inhibitors demonstrated improved event free survival in a curative setting in locoregionally advanced disease when given as neoadjuvant prior to surgery and with postoperative adjuvant therapy in addition to adjuvant with chemotherapy and radiation^14^. However, a clear improvement in overall survival has not been demonstrated. Increasingly, it has become clear that tumor-draining lymph nodes are critical to coordinating the immune response to the primary tumor, and concurrent ENI can abrogate the response to ICI. For example, cytotoxic radio-targeting of the tumor-draining lymph nodes in ENI disrupts dendritic cell-dependent, antigen-specific CD8-driven antitumor immunity, ceasing antitumor response^9,15^. To address this challenge, stereotactic body radiotherapy (SBRT) can deliver ablative high-beam radiation doses while minimizing the volume of irradiated normal tissue and sparing draining lymphatics, facilitating ICI response^16^. Data from clinical trials using tumor-targeted lymphatic-sparing SBRT in HNSCC patients are associated with increased local control and an immunostimulatory tumor microenvironment (TME), specifically when combined with systemic therapy agents such as immune checkpoint inhibitors (ICI)^17,18^. For example, when neoadjuvant SBRT is combined with a prescription immunotherapy anti-PD1 drug such as nivolumab, there is a major pathological response (mPR) rate of 60% in HPV-negative HNSCC as compared to the 7% seen with nivolumab alone, and a mPR rate of 100% in HPV-positive HNSCC as compared to 83% with SBRT alone^19^. The enhanced pathological response observed may be attributed in part to SBRT induced upregulation of PD-L1 expression in tumors initially PD-L1 negative, thereby rendering tumors more susceptible to anti-PD1/PD-L1 therapy^20^. Therefore, SBRT immunomodulation may sensitize otherwise unresponsive tumors to immune checkpoint inhibition. In addition, after SBRT and ICI therapy, the removal of regional lymphatics did not diminish the immune response, meaning this treatment can be used along with the current standard of lymphatic ablation when treating HNSCC^9^. However, and critically, although radiation increases tumor antigenicity and PD-1 blockade inhibits T cell exhaustion, tumors can modulate the surrounding microenvironment to escape therapy by upregulating *other* immunosuppressive markers^21^. CD47 is an immunosuppressive immune checkpoint receptor overexpressed by tumors to evade recognition by the immune system^22^. CD47 interacts with the ligand SIRPα found in myeloid and dendritic cells and conveys a “do not eat me” signal to inhibit tumor phagocytosis, thus blocking ingestion of tumor-specific antigens by professional APCs^22^. Although CD47 is a promising target for ICI, the utility of conventional CD47 blockade is compromised due to on-target, off-tumor phagocytic toxicities caused by extensive expression of CD47 on red blood cells and normal tissue^22^. Some researchers have identified routes around this limitation by using an engineered CD47 inhibitor with an inactive Fc region, evorpacept (ALX148), to avoid hematological toxicity^3^. These studies have found that the engineered anti-CD47 inhibitor promotes an antitumor immune response through methods such as increased macrophage phagocytosis, dendritic cell activation, and proinflammatory cytokine production while maintaining a favorable safety profile^1,3^. However, many of these studies do not investigate the lymphatic sparing immunoradiotherapy benefit when combined with anti-CD47 activity. For this study, we used an engineered CD47-blocking SIRPα fusion protein (ALX301) with an N297A mutation to minimize interaction with the target cell Fc receptor, decreasing off-tumor toxicity. We hypothesized that CD47 blockade would enhance PD-1 inhibitor and radiation tumor response in HNSCC by enhancing phagocytic tumor-specific antigen (TSA) presentation, resulting in improved immune-mediated tumor cell death via (enhanced cytotoxic T cell-mediated cell death) expansion of the TSA T cell repertoire and the immunopeptidome. We then placed these strategies in the context of a model for combinatorial immunoradiotherapy designed to spare injury to tumor draining lymphatics and thereby enhancing antitumor response.

## Results

To explore the clinical relevance of CD47 expression in HNSCC, we used patient data from the Cancer Genome Atlas (TCGA) dataset. Kaplan-Meier survival analysis showed that patients with higher CD47 expression had significantly lower disease-free survival than did those with lower expression, suggesting that CD47 may serve as a marker of poor prognosis (Figure 1A). These findings are consistent with other studies finding that lower expression of CD47 led to a significant increase in survivability^23^. To evaluate the impact of CD47 expression in preclinical HNSCC models, we employed previously characterized 4-NQO carcinogen-induced, 4MOSC1 murine oral squamous cell carcinoma cancer cell lines that can be employed in syngeneic, orthotopic models. This model displays a human tobacco-signature mutanome in addition to an immune infiltrate and ICI response pattern similar to that observed clinically in HPV-negative HNSCC, with partial response to anti-PD1 therapy^24^. To define the role of CD47 in this model, we created a CD47 knockout from our 4MOSC1 cell line to investigate the effect on tumor growth and the TME. We used a selective CRISPR antigen removal lentiviral vector system to ensure the elimination of Cas9 from the genome later, preventing immunogenicity^25^. Cas9 immunogenicity had been verified in Cas9-expressing 4MOSC1 buccal tumor-bearing mice (Figure 1B). Knockout of CD47 and Cas9 was verified as shown (Figure 1C). We observed that knocking CD47 out of 4MOSC1 sensitizes the cell line and results in complete tumor regression (Figure 1D). Using the CD47 KO Cas9 KO 4MOSC1 cell line, we further observed complete tumor regression in the knockout groups, whereas the parent 4MOSC1 cell line tumors continued to grow (Figure 1E).

**Figure 1.**
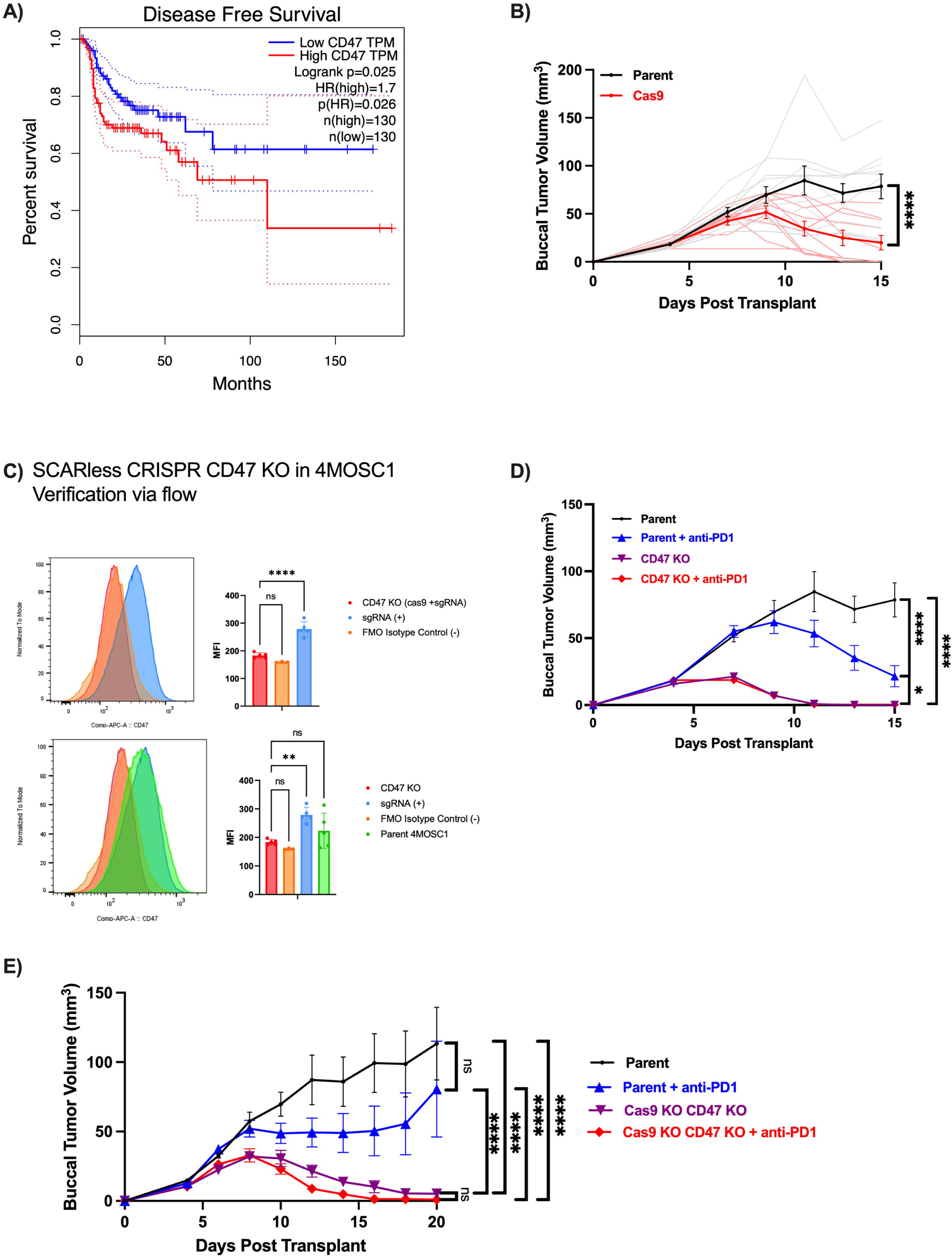
**a.** Kaplan Meier survival analysis of TCGA HNSC data set based on CD47 expression ^26^. A quartile cut-off was used and survival analysis was performed using log-rank tests. **b.** Representative tumor growth kinetics of mice with tumors of the parent 4MOSC1 cell line vs. tumors with Cas9 expressing 4MOSC1 cell line. **c.** Flow cytometry results demonstrating knockout of CD47 and Cas9 from the 4MOSC1 parent cell line. FMO isotype controls are used to determine the cut-off point between background fluorescence and positive populations in multi-color immunofluorescent experiments. 4MOSC1 cells were first transduced with pSCAR_Cas9 GFP-expressing vector, infected with pSCAR_sgRNA targeting CD47, then cultured for 10 days to allow Cas9 genome editing and removal of CD47, and later the IDLV-Cre vector expressing Cre recombinase was transduced to excise the loxP site containing Cas9. **d.** Representative tumor growth kinetics of mice with 4MOSC1 parent tumors vs. 4MOSC1 parent tumors treated with anti-PD1 vs. 4MOSC1 CD47 knockout tumors vs. 4MOSC1 CD47 knockout tumors treated with anti-PD1. **e.** Representative tumor growth kinetics of mice with 4MOSC1 parent tumors vs. 4MOSC1 parent tumors treated with anti-PD1 vs. 4MOSC1 CD47 Cas9 knockout tumors vs. 4MOSC1 CD47 Cas9 knockout tumors treated with anti-PD1 are shown.

Based on the results obtained from CD47 4MOSC1 knockout models, we chose to observe how these findings would translate through CD47 inhibition obtained through pharmacologic CD47 blockade using ALX301. ALX301 is an engineered CD47-blocking SIRPα fusion protein with an N297A mutation which minimizes interaction with the myeloid cell Fc receptor, decreasing off-tumor toxicity^22^. To explore the potential role of pharmacologic anti-CD47 treatment, we defined response to ALX301 in models employing the anti-PD1 partially responsive 4MOSC1 model as well as an anti-PD1 resistant 4MOSC2 mode^24^. Mice transplanted with the 4MOSC1 or 4MOSC2 cell line were treated with either anti-CD47 (30mg/kg every 4 days starting on day 6), anti-PD1 (10mg/kg on days 6 and 8), or both and monitored for tumor growth/regression. Combination treatment of anti-CD47 with anti-PD1 led to a complete tumor regression in the 4MOSC1 model (Figure 2A), but only a partial tumor regression in the 4MOSC2 cell line (Figure 2B). To overcome the partial tumor regression observed in the 4MOSC2 cell line, a triple regimen of anti-CD47, anti-PD1, and tumor-directed radiation therapy (tdRT) was designed (Figure 2C). The addition of tdRT therapy was based on the immunomodulatory effects observed after SBRT was implemented in both preclinical and clinical settings. A one-time dose of 4 Gy was chosen for tdRT based on results that provided evidence that this dose of tdRT was not cytotoxic but still enhanced the percentage of CD8+ T cells and dendritic cell trafficking to the tumor. With SBRT, there was an enrichment of tumor-specific T cells and CD8+ cytotoxic T cells, indicating an enhancement of an antitumor response^18^. Furthermore, we chose to deliver the tdRT before ICI administration due to evidence that delivering irradiation treatment directly to the gross tumor and sparing the cervical lymphatics can improve the systemic response that anti-PD1 and other immunotherapies provide^15, 19^. Figure 2C shows that the introduction of tdRT to the treatment allows further tumor regression in mice with the 4MOSC2 tumor model. Treatment with anti-PD1, anti-CD47, or a combination, along with tdRT increased long-term survivability (n=10 for each arm, control vs. anti-PD1+tdRT p=0.0505, control vs. anti-PD1+anti-CD47 p=0.0114*, control vs. anti-PD1+anti-CD47+tdRT p=0.016*; Supplemental Figure 1C). Of note, enhanced long-term survivability was associated with anti-CD47 as a part of treatment, illustrating the benefits of anti-CD47 in combination therapy. Based on these findings, there is evidence that an anti-CD47 ICI in combination with either an anti-PD1 ICI or 4 Gy tdRT can overcome an immunosuppressive TME.

**Figure 2.**
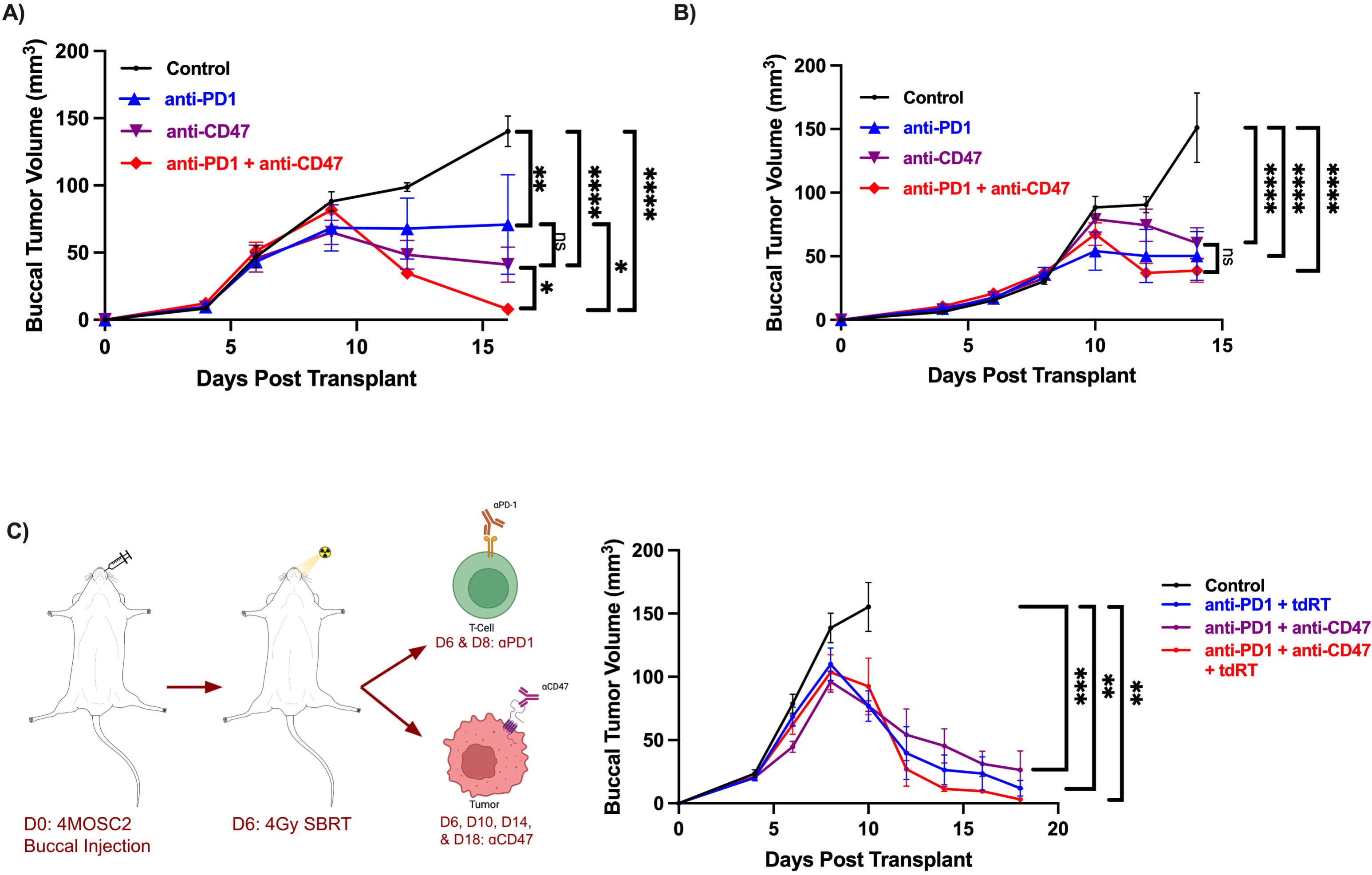
**a.** Representative tumor growth kinetics of mice with 4MOSC1 tumors (control) vs. 4MOSC1 tumors treated with anti-PD1 vs. 4MOSC1 tumors treated with anti-CD47 (ALX301) vs. 4MOSC1 treated with both anti-CD47 and anti-PD1 are shown. **b.** Representative tumor growth kinetics of mice with 4MOSC2 tumors (control) vs. 4MOSC2 tumors treated with anti-PD1 vs. 4MOSC2 tumors treated with anti-CD47 vs. 4MOSC2 treated with both anti-CD47 and anti-PD1 are shown. **c. (Left)** Graphic representation of the experimental design for the triple regimen of anti-PD1, anti-CD47, and tdRT. The 4MOSC2 cell line will be transplanted into the buccal mucosa on day 0 for all mice. Mice who receive tdRT will receive 4Gy of tdRT on day 6. Mice who receive anti-PD1 treatment will receive it on day 6 and day 8, and mice who receive anti-CD47 treatment will receive it on day 6, day 10, day 14, and day 18. **(Right)** Representative tumor growth kinetics of the experimental design shown to the left with 4MOSC2 (control) vs. 4MOSC2 tumors treated with anti-PD1 and tdRT vs. 4MOSC2 tumors treated with anti-PD1 and anti-CD47 vs. 4MOSC2 treated with anti-PD1, anti-CD47, and tdRT.

To better understand the mechanism by which anti-CD47 ICI can provide antitumor immunity, we characterized tumor, sentinel lymph nodes (SLN), and central lymph nodes (CLN) on a cellular level. We first transplanted mice with 4MOSC1 tumors and treated one group with the vehicle and the other with 3 doses of ALX301, where dosages were administered every 4 days starting on day 6 at a concentration of 30mg/kg. On day 15 post-transplant, the SLN and CLN were mapped using lymphazurin to visually confirm the tumor draining sentinel lymph node (SLN) and then harvested along with the tumor (Figure 3A). Anti-CD47 inhibited tumor growth (Figure 3B), and we noted upregulation in MHC II in CD11c+ MHC II+ dendritic cells (DC) in the tumor (Figure 3C). This suggests that anti-CD47 enhanced immune activation through DCs being able to more effectively present antigens to T cells, leading to a stronger T cell-mediated tumor regression. We also noted an increase in CD86 expression on activated DCs in the tumor, SLN, and CLN (Figure 3D), suggesting a co-stimulatory signal on activated CD4+ T cells that works synergistically with MHC II to activate and prime T cells for an anti-tumor response. Taken together, these results demonstrate that anti-CD47 treatment as a standalone therapy elicits an enhanced immune response through upregulation of MHC II and CD86 on DCs allowing for greater antigen presentation, T-cell activation, and T-cell expansion. Of note, the effect of anti-CD47 on other immune parameters, including the count of CD8 T-cells in the tumor, CD86 MFI in F4/80+ macrophages, and percent CCR7+ activated DCs were not statistically demonstrated.

**Figure 3.**
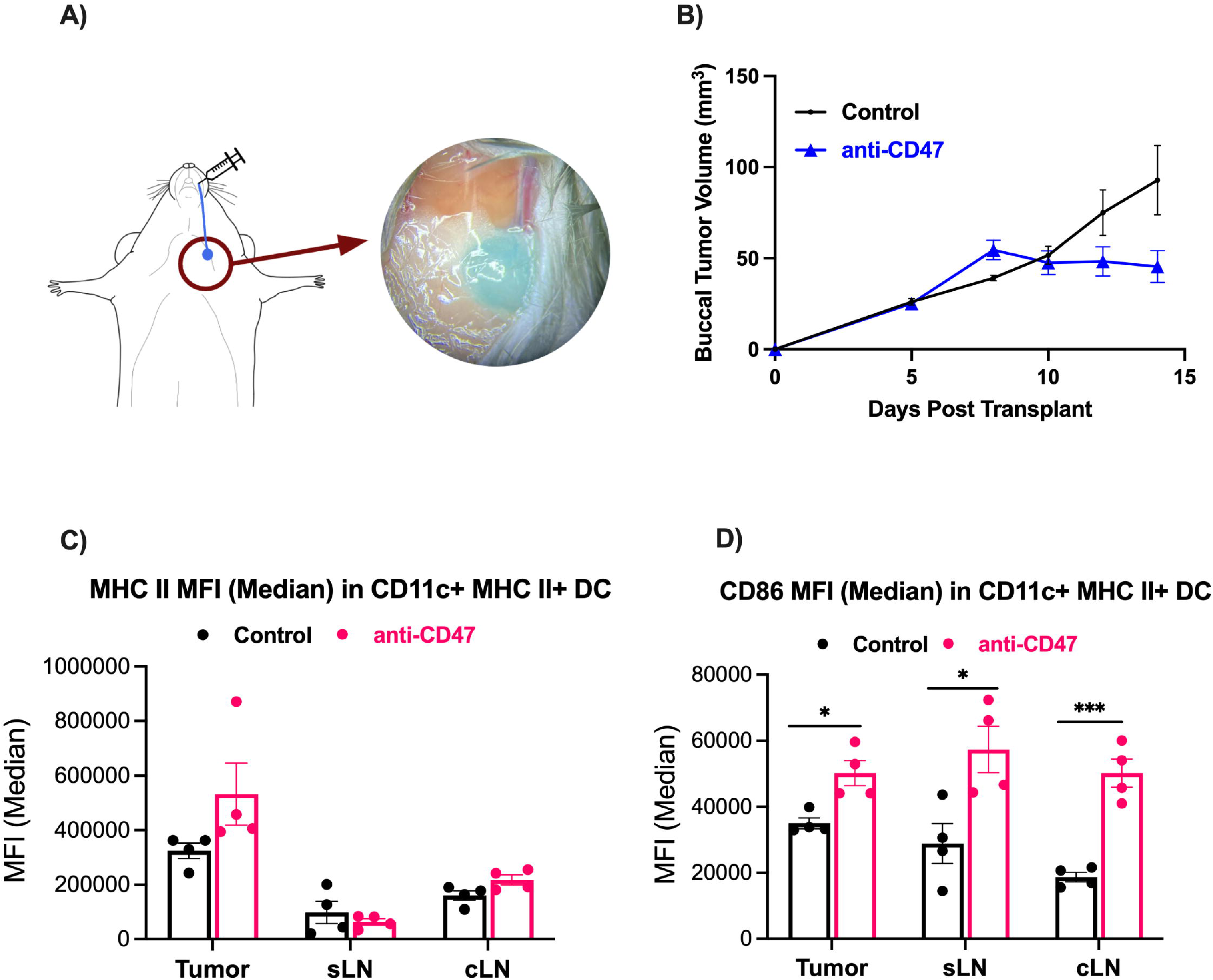
Anti-CD47 therapy reduces tumor burden and enhances immune activation. Mice were injected with 4MOSC1 cells (1×10^6^ cells) in the buccal mucosa, and on days 6, 10, and 14 were given 30mg/kg of ALX301 (anti-CD47) before being assessed on Day 15. **a.** Process of lymphatic mapping of the cervical draining lymph node in the 4MOSC1 mouse model. Lymphazurin is injected into the buccal mucosa and the draining lymph node is marked blue. **b.** Representative tumor growth kinetics of mice with 4MOSC1 tumors (control) vs. 4MOSC1 tumors treated with anti-CD47. **c.** In the tumors of the anti-CD47 treated group, there was an upregulation in MHC II in CD11c+ MHC II+ dendritic cells. **d.** In the tumors, SLNs, and CLNs of the anti-CD47 treated group, there was an upregulation in CD86 in CD11c+ MHC II+ dendritic cells.

To explore how anti-CD47 can be combined with other therapeutic modalities, we employed our ICI resistant 4MOSC2 model and treated it with anti-CD47, anti-PD1, and tdRT in a fashion similar to that depicted in Figure 2C. (Supplemental Figure 1E). Tumors of the mice in all treatment groups demonstrated upregulation of CD69 in CD8+ T cells (Figure 4), consistent with the effects of anti-PD1 in upregulating T cell activation within the TME as well as in increasing T cell retention in the tumor for a sustained local immune response^27^. Additionally, we did not demonstrate changes in CD69 MFI in natural killer (NK) cells, macrophage recruitment (F4/80+ Mac% in CD45+), and CD86 MFI in F4/80+ macrophages, consistent with a primary effect of treatment on end effector CD8+ T cells.

**Figure 4.**
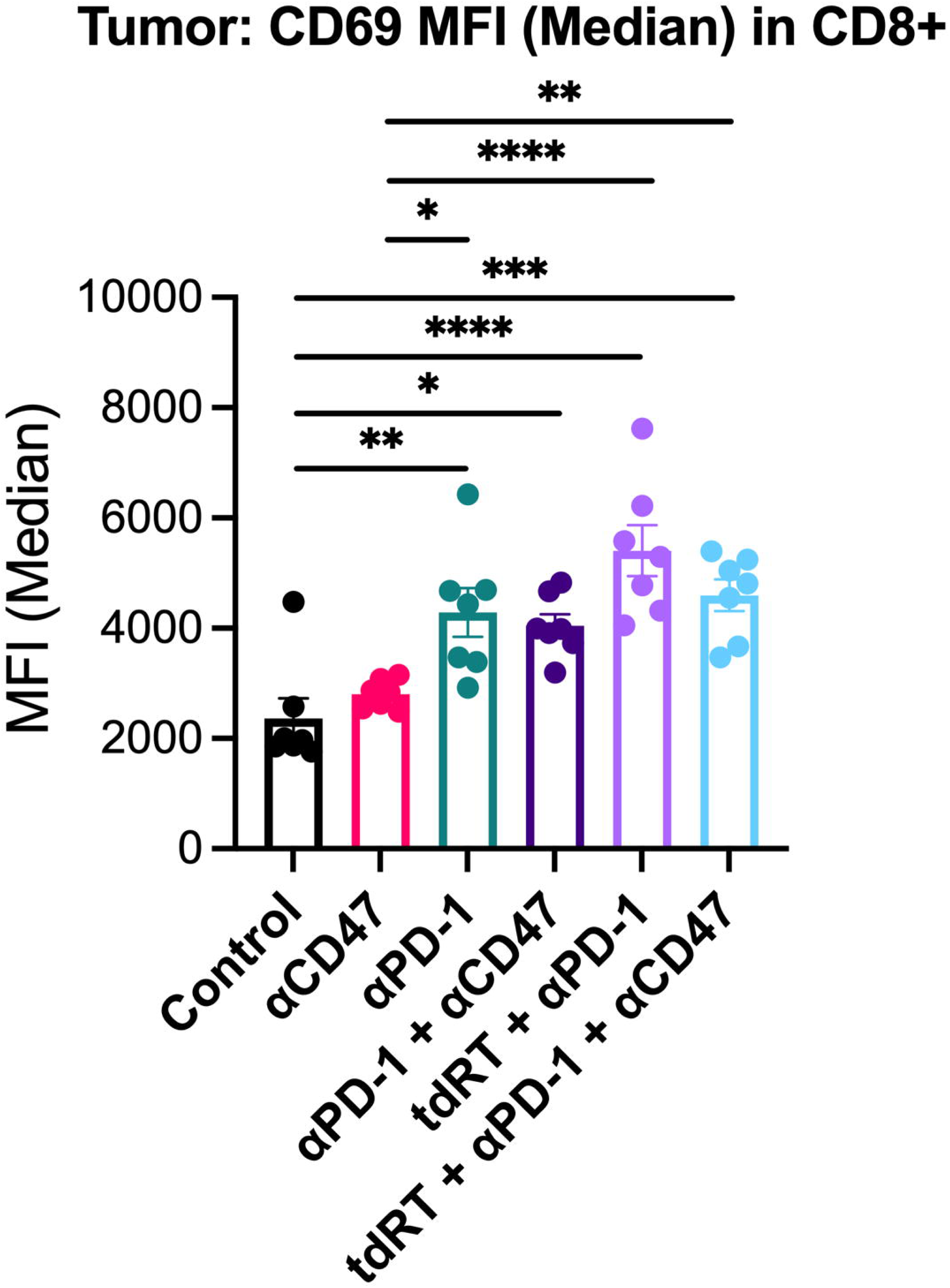
Mice were injected with 4MOSC2 cells (0.5×10^6^ cells) in the buccal mucosa before being assessed on Day 9. Mice in the anti-PD1 and all combination treatment groups observed an upregulation of CD69 in CD8+ T cells in the tumor.

To specifically assess how the treatment of anti-CD47, anti-PD1, and tdRT affects the T-cell response, we performed TCR sequencing on the tumors and SLNs of mice treated above (Figures 3 and 4). TCRa and TCRb clonality analysis demonstrates that in both 4MOSC1 and 4MOSC2 models, there is a significant enhancement in clonal expansion in anti-CD47 treated mice, with the one exception of 4MOSC2 anti-PD1+tdRT tumors (which may be due to a large therapeutic response to the combination therapy of anti-PD1 and tdRT). We also defined overlap of TCRa and TCRb clonotypes between the tumors and SLN in the 4MOSC1 control vs. anti-CD47 experiment (Figure 5B and 5C) showing an increase in common TCR clones in anti-CD47 treated tumors. Interestingly, in 4MOSC2 models, we saw greater shared clonality between the tumors and SLNs in mice treated with anti-CD47 (Figure 5D, 5E, 6). The increase in shared clonality between the tumors and SLNs indicated that the T cells that were activated and expanded within SLN in response to anti-CD47 had migrated to the TME, potentially further amplifying the immune response. This also reinforces the key role of antigen exposed, migratory DCs to the SLN and draining lymph nodes in antigen specific activation of tumor-infiltrating lymphocytes (TILs) and response to tumor-associated antigens. Ultimately, these results suggest that anti-CD47 plays a critical role in the expansion and migration of specific T-cell clonotypes in response to tumor-associated antigens, and that anti-CD47 can specifically enhance anti-tumor specific T cell repertoire.

**Figure 5.**
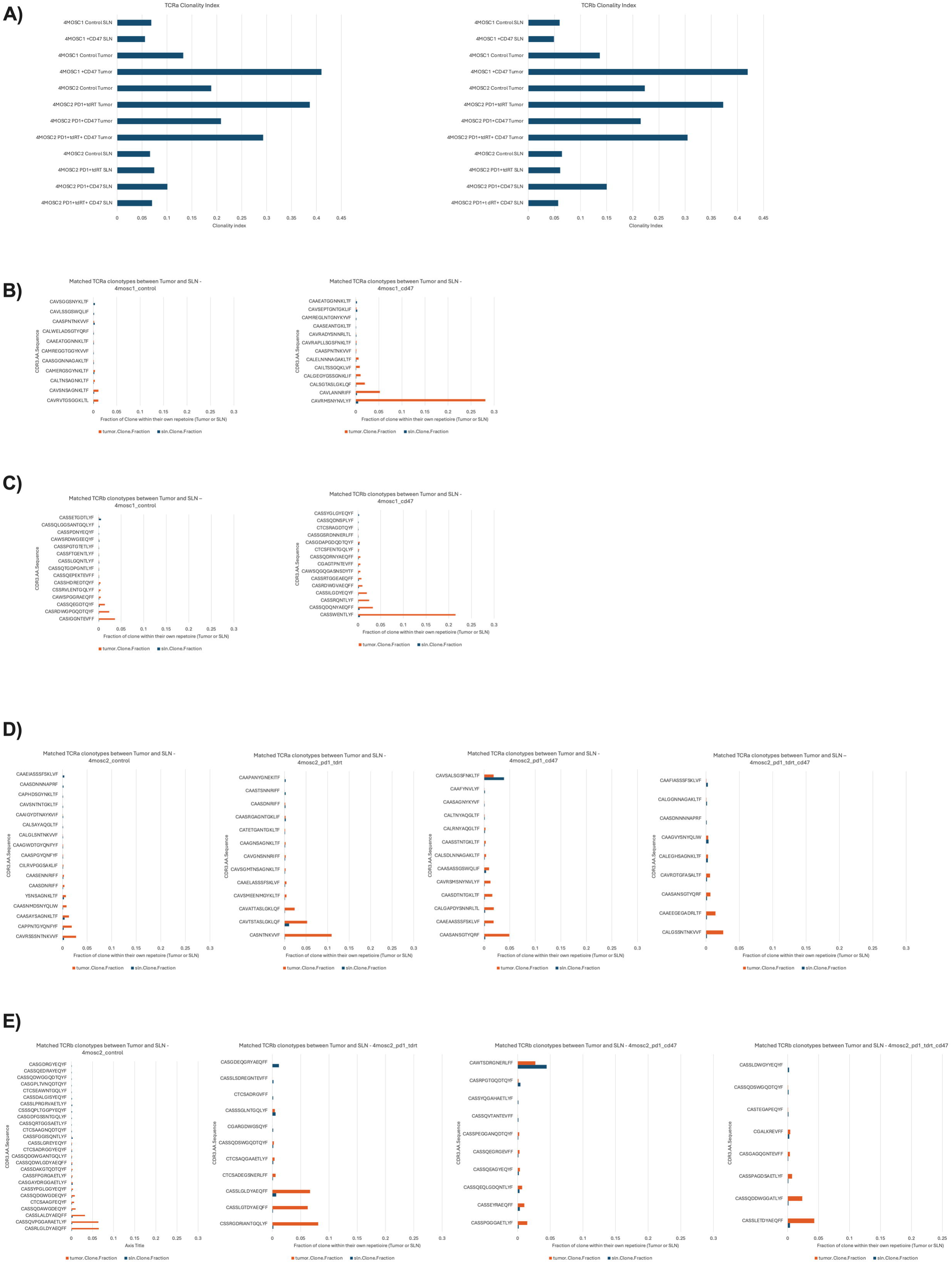
TCR sequencing of 1 tumor and 1 SLN from each treatment group in Figure 3 (4MOSC1 control vs. anti-CD47) and Figure 5 (4MOSC2 control vs. anti-PD1+anti-CD47 vs. anti-PD1+tdRT vs. anti-PD1+anti-CD47+tdRT). TCR sequencing demonstrates that anti-CD47 treatment leads to upregulation of clonal expansion within the tumor and shared clonality between the tumors and SLNs. **a.** TCRa and TCRb Clonality Index of the tumors and SLNs of the different models and treatment groups. **b.** Matched TCRa clonotype between tumors and SLNs of the 4MOSC1 model. **c.** Matched TCRb clonotype between tumors and SLNs of the 4MOSC1 model. **d.** Matched TCRa clonotype between tumors and SLNs of the 4MOSC2 model. **e.** Matched TCRb clonotype between tumors and SLNs of the 4MOSC2 model.

**Figure 6.**
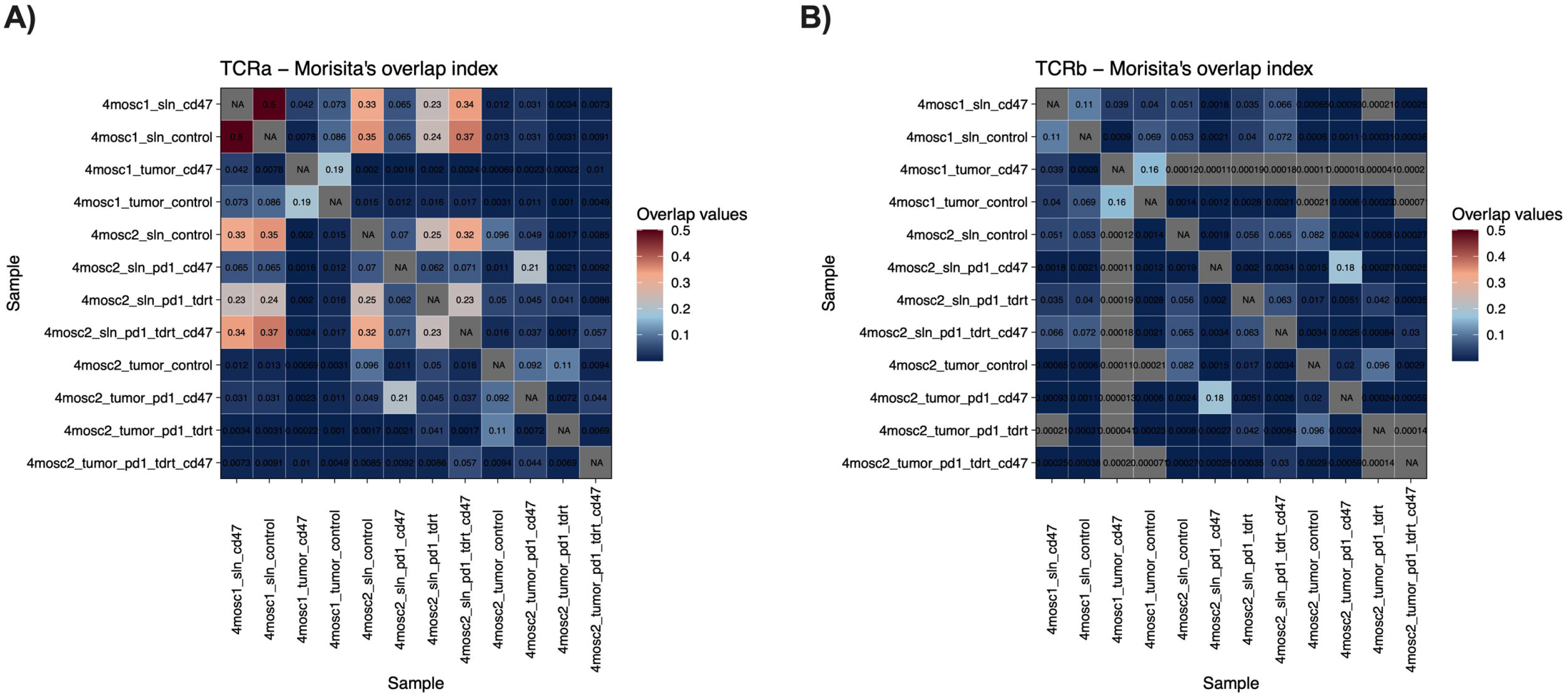
Morisita’s Overlap Index of the TCR-sequencing data observed in Figure 5 demonstrating increased shared clonality between tumors and SLNs of anti-CD47 treated groups. **a.** TCRa clonality for both 4MOSC1 and 4MOSC2 models. **b.** TCRb clonality for both 4MOSC1 and 4MOSC2 models.

## Discussion

The rise of ICI therapy such as immunotherapy targeting PD1/PD-L1 has shown promise as an effective treatment with low toxicity^28^. However, despite the overall increase in survivability in patients who receive PD1 ICI treatments, recent studies demonstrate that most patients with HNSCC may not respond to PD1 ICI therapy or will develop resistance to the immunotherapy after some time^29^. In fact, in the Javelin and Head Neck 100 trials, it was demonstrated that avelumab failed to improve the progression-free survival and overall survival of HNSCC compared to other types of cancers, such as non-small cell lung cancer, which saw improved outcomes^30^. These data suggest a need for other types of therapies that are just as or more effective whether alone or in combination with ICIs^30^. Clinical trials have also demonstrated that sequencing of therapies are important, for example, when anti-PD1 is delivered before chemoradiotherapy (CRT), there is an abrogated immune response due to radiosensitive CD8+ T cells, whereas if anti-PD1 is delivered after CRT, there is better control of the radiated tumor^30^. However, not only was the sequencing of the combined therapies found to be important, but also whether the tumor-draining lymph nodes were included in the radiation fields. In clinical treatments of HNSCC, current evidence shows that the antitumor immune response derived from PD-1/PD-L1 inhibitors results from the CD8+ T cells within the tumor-draining lymph nodes^31^. When these lymph nodes are included in the radiation field, it leads to a depletion of the antitumor CD8+ T cells, therefore inhibiting the host’s antitumor immunity^30^. Given these considerations, we designed a new combinatory therapy for our study. As CD47, another immune checkpoint located upstream of PD-1 in the adaptive tumor immune response pathway, is highly expressed in many tumors and is targetable, this presents a good candidate for combination treatment along with PD1 ICIs. Moreover, these therapies paired with SBRT could target the tumor without exposing the lymph nodes to the elective radiation volume, further optimizing treatment.

CD47 has long been studied as a potential target of immunotherapy in cancer treatments. CD47 was first identified as a tumor antigen in the 1980s, and since then, has been found to be expressed on multiple human tumor types and at increased levels in tumors compared to their normal cell counterparts^2^. Current evidence supports the idea that the over-expression of CD47 allows tumors to evade the immune system by binding to SIRPα, allowing tumor cells to express the “Don’t Eat Me” signal to evade phagocytosis, promoting tumor survival and growth^2^. Based on these conclusions, a CD47-blocking myeloid checkpoint inhibitor, evorpacept (ALX148), was developed to promote phagocytosis of tumor cells. It was found to have a favorable safety profile in clinical studies^3^. Additionally, in preclinical models, evorpacept combined with anti-PD1 led to favorable outcomes such as increased tumor growth inhibition and extended survivability^1^.

While phase 2 clinical trials investigating evorpacept, pembrolizumab, and chemotherapy combination therapy for locally advanced HNSCC failed to meet primary endpoints, these findings suggest that additional or alternative therapeutic partners may be necessary for the synergistic potential of CD47 and PD1 inhibition (NCT05787639 and NCT04675333)^32^. Based on these results, we chose to explore a similar anti-CD47 ICI in combination with other therapeutics while avoiding lymph node ablation.

In our study, after confirming the prognostic association of CD47 expression with prognosis in HNSCC, we first confirmed the effectiveness of CD47 as a viable target for immunotherapy. After knocking CD47 out of the 4MOSC1 genome, we demonstrated that the CD47 knockout tumor cells were significantly more susceptible to anti-tumor immune response than the parent tumor cells even when the parent tumor was treated with an anti-PD1 ICI. After establishing that CD47 is a viable target for HNSCC therapeutics, we used an engineered CD47-blocking SIRPα fusion protein (ALX301) with an N297A mutation to treat 4MOSC1/4MOSC2 HNSCC in a murine model and demonstrated its effectiveness as a standalone therapeutic as well as in combination with an anti-PD1 ICI. In our 4MOSC1 tumor model, we found that treating mice with either anti-PD1 or anti-CD47 provided better results than the control group with similar response. However, by combining anti-PD1 and anti-CD47 in combination, we were able to achieve complete regression of 4MOSC1 tumors. Additionally, in our PD-1 resistant 4MOSC2 model, using both anti-PD1 and anti-CD47 showed significant response, and the addition of anti-CD47 enhanced survival. Interrogation of the immune microenvironment showed that in anti-CD47 treated 4MOSC1 tumors there was an increase in activated dendritic cells in tumors and draining sentinel nodes, accompanied by an increase in activated T cells in the TME. When we combined anti-CD47 with anti-PD1 and tdRT as a triple regimen against our aggressive 4MOSC2 model, we also saw increased activation and retention of CD8+ T cells. After performing TCR sequencing on these models, we observed with interest that, when tumors were treated with anti-CD47, there was an increase in total T-cell clonal expansion and in the fraction of clones within their own repertoire as well as an increase in shared clonality between the tumor and SLN associated with enhanced antigen specific adaptive immune response. This is consistent with an enhanced clonal T cell response generated in draining sentinel lymph nodes with migration of T cells from the draining SLN to TME. Ultimately, these data support our hypothesis that anti-CD47 increases trafficking and migration of APCs, activation of T-cells, and priming and expansion of T cells against tumor antigens.

These findings support the rationale for CD47 inhibition in combination with current immunomodulatory treatments such as anti-PD1 ICIs and SBRT therapies. HPV-negative HNSCC is associated with a poor prognosis, so it is crucial to define novel, effective therapeutic strategies that can enhance response^5^. By demonstrating that anti-PD1 and radiotherapy can be further optimized in combination with anti-CD47, we have provided a basis for rational combination of these therapies to enhance tumor response.

However, although we have discovered promising results in our treatment of the syngeneic HPV-negative HNSCC, animal models are always limited: they do not capture the full complexity of the disease in a clinical setting. Moreover, due to the short duration of treatment of the 4MSOC tumors, we were unable to evaluate the toxicity of the regimen. In HNSCC, the mechanism of CD47 inhibition, and the efficacy with which it impacts tumor progression, is still largely unknown. Previous studies have demonstrated that CD47 is overexpressed in HNSCC and that its inhibition as a standalone therapy has led to the stimulation of effector T cells and a decrease in suppressive immune cells^33^. However, as we have demonstrated in our own study, the benefits of CD47 inhibition as a combination therapy reveal its promise as a future HNSCC treatment. CD47 inhibition is not limited to combination with anti-PD1 and tdRT: it has shown great results synergistically with other therapies as well. One of the most prominent cancer therapeutics, tumor targeted antibodies, has been shown to be enhanced when combined with anti-CD47, notably because of the enhanced phagocytosis from anti-CD47 in combination with antibody-dependent activation of macrophages through the Fc-gamma receptor^34^. These findings, along with our own, illustrate the multiple advantages of introducing a CD47 inhibition into the range of current practices as a method of enhancing antitumor activity.

As clinical studies investigating anti-CD47 therapies progress, leveraging syngeneic HNSCC models to derive rational sequencing and design of combination therapies can help accelerate and optimize clinical trial design. In addition, mechanistic understanding of the ways in which CD47 inhibition interacts with current therapies, including those that affect draining lymphatic basins, can help to synergize CD47 inhibition with current standard of care therapy^15^. These data can help further optimize how we enhance antitumor activity while reducing tumor evasion mechanisms in both HPV-positive and negative HNSCC.

## Methods

All the animal studies were approved by the University of California San Diego (UCSD) Institutional Animal Care and Use Committee (IACUC, protocol #S16200); all experiments adhere with all relevant ethical regulations for animal testing and research.

### Cell lines and tissue culture

The 4MOSC (4MOSC1, 4MOSC2, 4MOSC1 CD47 KO, 4MOSC1 Cas9 KO, 4MOSC1 CD47 KO Cas9 KO) syngeneic mouse HNSCC cells harboring a human tobacco-related mutanome and genomic landscape were developed and described for use in immunotherapy studies in a prior report^24^. 4MOSC cells were cultured in Defined Keratinocyte-SFM medium supplemented with EGF Recombinant Mouse Protein (5 ng/ml), Cholera Toxin (50 pM), and 1% antibiotic/antimycotic solution. 293T cells (ATCC CRL-3216) were cultured in Dulbecco’s Modified Eagle’s Medium (DMEM) supplemented with 10% fetal bovine serum, 2 mM L-glutamine (ATCC 30-2214) and 1% antibiotic/antimycotic solution. All cells were cultured at 37L°C in the presence of 5% CO_2_.

### Cloning of pLenti-eGFP-LucOS

The full-length coding sequence of LUC-OS flanked by attbB1/2 recombination site was amplified from the Lenti-LucOS (22777) using the LUC-OS-F (5′-GGGGACAAGTTTGTACAAAAAGCAGGCTTAATGGAAGACGCCAAAAACATA-3′) and LUC-OS-R (5′-GGGG ACCACTTTGTACAAGAAAGCTGGGTTTTACAAGTCCTCttCAGAAAT-3′) primer. The purified PCR product was incorporated into the pDONR221 vector via a BP Reaction and subsequently introduced into the pLenti-CMV-GFP-DEST (19732) through an LR reaction.

### Generation of stable GFP-Luc and eGFP-LucOS expressing 4MOSC1 and 4MOSC2 cell line

For lentivirus production, 293T cells were plated in a poly-D-lysine–coated 15-cm dish and, 16Lh later, transfected with 30Lmg pHAGE PGK-GFP-IRES-LUC-W or pLenti-eGFP-LucOS, 3Lmg VSV-G, 1.5 µg Tat1b, 1.5 µg Rev1b, and 1.5 µg Gag/Pol using 25.2 µL P3000 reagent and 25.2 µL of Lipofectamine 3000 transfection reagent, and media was refreshed 16Lh post-transfection. At 48 and 72 h, virus-containing media was collected, filtered througfh a low protein binding filter unit (PVDF, 0.45 µm, Sigma-Aldrich), and stored at 4 °C for up to 5 days prior to use. Lentivirus suspension was concentrated using Lenti-X concentrator per manufacturer standardized protocol (Takara Bio). Subsequently, 4MOSC1/2 cells were plated in a collagen-coated 6-well plate. At 16 h, seeded cells were transduced using 200 µL of concentrated virus in 2 mL keratinocyte-defined serum-free media and 4 µg/mL polybrene, and the plate was immediately centrifuged for 15 min at 450×g. GFP expression was validated by fluorescent microscopy and flow cytometry. Transduced 4MOSC1/2 cells were sorted by FACS for viability and GFP-positivity using a FACS-Aria Cell Sorter (BD Biosciences).

### Generation CD47 knockout 4MOSC1 cell line using selective CRISPR antigen removal lentiviral vector system

Lentiviral vectors pSCAR_Cas9, IDLV-Cre, and pSCAR_sgRNA (Cd47 sgRNA: CCACATTACGGACGATGCAA) were generated and obtained from a previous study^25^. 4MOSC1 cell line was first infected with pSCAR_Cas9 in media containing polybrene (Millipore Sigma #107689); after 48 hours they were selected with blasticidin (InvivoGen #ant-bl-1) for 7 days. Cas9 expression by 4MOSC1 was verified via western blot (Supplemental Figure 1B). Cas9-expressing cells were then infected with lentivirus pSCAR_sgRNA and after 48 hours, they underwent selection with blasticidin for 3 days. Following pSCAR_sgRNA transduction, cells were cultured in blasticidin for a total of 10 days to allow genome editing while maintaining high expression of SCAR vectors. At the end of 10 days, cells were infected with IDLV-Cre in media containing polybrene and blasticidin. Following IDLV-Cre transduction (by the same process as other lentivirus infections), cells were sorted by FACS for viability and GFP and mKatie negativity using FACS-Aria Cell Sorter (BD Biosciences).

### In vivo mouse models and analysis

All the animal studies using HNSCC orthotropic implantation studies were approved by the UCSD Institutional Animal Care and Use Committee (IACUC), with Animal Study Proposal (ASP) protocol #S16200; and all experiments adhere to all relevant ethical regulations for animal testing and research. All mice were obtained from Jackson Laboratories (San Diego, CA). Mice at UCSD Moores Cancer Center are housed in individually ventilated and micro-isolator cages supplied with acidified water and fed 5053 Irradiated Picolab Rodent Diet 20. The temperature for laboratory mice in this facility is mandated to be between 18 and 23 °C with 40–60% humidity. The vivarium is maintained in a 12-h light/dark cycle. All personnel were required to wear scrubs and/or lab coat, mask, hair net, dedicated shoes, and disposable gloves upon entering the animal rooms. WT C57Bl/6 mice were obtained from Jackson Laboratories (San Diego, CA).

### Orthotopic tumor modeling

For orthotopic implantation, 4MOSC1 cells were transplanted (1 million per tumor) into the oral cavity of female C57Bl/6 mice (4–6 weeks of age) to the buccal mucosa. 4MOSC2 cells were transplanted (0.5 million per tumor) into the buccal mucosa of the oral cavity of female C57Bl/6 mice (4–6 weeks of age). For drug treatment, the mice were treated by intraperitoneal injection (ip) of PD-1 antibody or CD47 antibody. The mice were sacrificed at the indicated timepoints (or when mice succumbed to disease—buccal tumors >1 cm in greatest dimension or ulcerated— as determined by the ASP guidelines) and tissues were isolated for histological and immunohistochemical evaluation or flow cytometric analysis. The maximum tumor size/burden permitted in accordance with our institutional review board was not exceeded.

### Lymphatics mapping

Lymphatic mapping was done by injecting lymphazurin into the buccal mucosa and following the flow of the blue dye to the cervical draining lymph node. The lymphazurin was reconstituted using isosulfan blue (HY-107967).

### Radiation

For the tumor-directed Radiation Therapy (tdRT), mice were treated with one dose of 4Gy stereotactic body radiotherapy (SBRT) focused on the buccal mucosa containing the tumor. tdRT was done on day 6 post transplant of the tumor prior to ICI treatments.

### Preparation of ICI treatments

All anti-PD1 treatments were prepared at a dose of 10mg/kg and mice were treated on days 6 and 8. All anti-CD47 treatments were prepared at a dose of 30mg/kg and mice were treated on days 6, 10, 14, and 18 except for flow cytometry experiments where takedown occurred earlier. The vehicle used for the control was PBS.

### Flow Cytometry Analysis

Immunophenotyping of tumor-bearing mice was performed according to the STAR protocol with some modifications^35^. Briefly, tumor tissue, sentinel lymph nodes, and non-sentinel lymph nodes were mapped and harvested using lymphazurin. To obtain a single-cell suspension, tissues were passed through 70 μm cell strainers and treated with ACK buffer for red blood cell lysis.

The single-cell suspension was then stained with Zombie NIR™ Fixable Viability Kit (BioLegend). After Fc blocking with TruStain FcX™ (anti-mouse CD16/32 antibody) (BioLegend), the cells were stained with fluorochrome-conjugated antibodies in Brilliant Stain Buffer (BD Biosciences). The cells were fixed with 2% paraformaldehyde and analyzed on Agilent NovoCyte Advanteon (Agilent Technologies) with standard lasers and optical filters. Data was analyzed using FlowJo (FlowJo, LLC). The following fluorochrome-conjugated antibodies (clone, dilution) were used in this study: CD45.2 (104, 1:200), CD8α (53-6.7, 1:200), NK1.1 (PK136, 1:100), CD11c (N418, 1:200), F4/80 (BM8, 1:100), CD197/CCR7 (4B12, 1:50), CD69 (H1.2F3, 1:100), CD86 (GL-1, 1:100) from BioLegend, and I-A/I-E (2G9, 1:500) from BD Biosciences.

### TCR Sequencing

TCR sequencing was prepared by harvesting one tumor and one SLN from one mouse of each treatment group. Immediately after harvesting, RNA was prepared using QIAGEN RNA extraction kit. RNA samples were sent to CD Genomics for sequencing.

### Statistics

Data analysis was performed using GraphPad Prism 10 for MacOS. Differences in experimental groups were analyzed using independent t-tests, one-way ANOVA with multiple comparisons, or two-way ANOVA with multiple comparisons. Survival Analysis was performed using the Kaplan-Meier method and Survival Analysis. All data is reported as mean ±LSEM.

## Acknowledgements

We thank ALX Oncology for providing the ALX301 (anti-CD47) reagent used in this study. Cartoon renderings were created with the BioRender online platform (BioRender.com).

## Supporting Information Captions

**Supplemental Figure 1.**
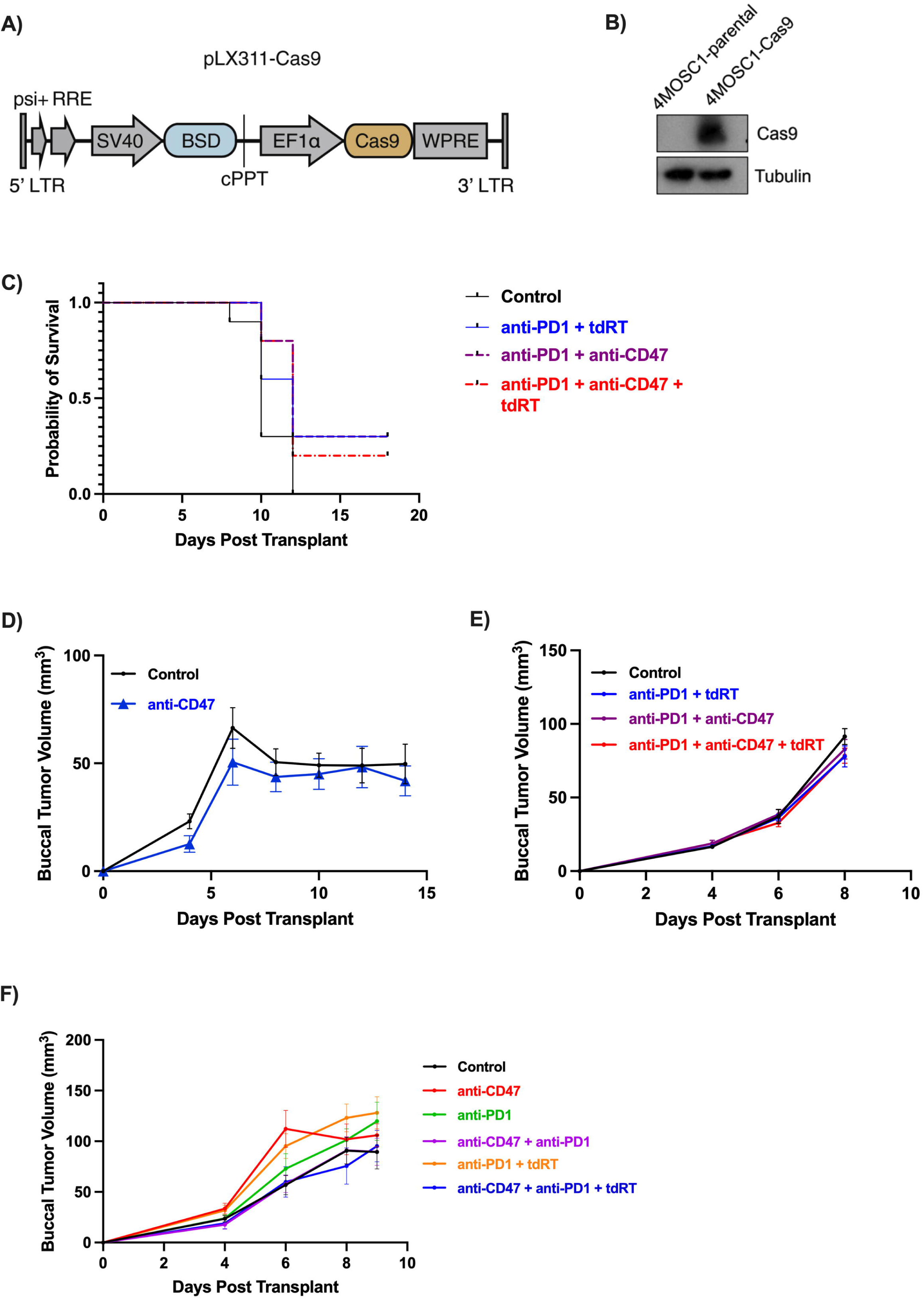
**a.** pSCAR_sgRNA (Cd47 sgRNA: CCACATTACGGACGATGCAA) used in the knockout of the Cas9 and CD47 represented in Figure 1C^25^. **b.** Verification of Cas9 expression in 4MOSC1 using western blot. **c**. Kaplan Meier curve demonstrating the survivability from the experiment depicted in Figure 2C (n=10 for each arm, Control vs anti-PD1+tdRT p=0.0505, Control vs anti-PD1+anti-CD47 p=0.0114*, Control vs anti-PD1+anti-CD47+tdRT p=0.016*) where with the treatment of anti-CD47, there is a statistically significant increase in survivability. **d.** Representative tumor growth kinetics of mice with 4MOSC1 tumors treated with vehicle (control) vs 4MOSC1 tumors treated with anti-CD47. **e.** Representative tumor growth kinetics of mice with 4MOSC2 tumors treated with vehicle (control), anti-CD47, anti-PD1, anti-CD47+anti-PD1, anti-PD1+tdRT, and anti-CD47+anti-PD1+tdRT.

